# Generation and therapeutic efficacy of the novel Fc-modified cross-neutralizing human monoclonal antibodies against dengue virus

**DOI:** 10.1101/2023.05.25.542225

**Authors:** Subenya Injampa, Surachet Benjathummarak, Sujitra Keadsanti, Wilarat Puangmanee, Atsushi Yamanaka, Tadahiro Sasaki, Kazuyoshi Ikuta, Pongrama Ramasoota, Pannamthip Pitaksajjakul

## Abstract

**Background:** Dengue disease is a mosquito-borne infection caused by four dengue virus serotypes (DENV1–4). Secondary infections can produce flavivirus cross-reactive antibodies at sub-neutralizing levels. This phenomenon can significantly increase the severity of secondary infections via antibody-dependent enhancement (ADE). ADE activity is associated with a high risk of viral infection in immune effector cells, triggering cytokine cascade and activating the complement system, which lead to severe symptoms. Despite extensive studies, therapeutic antibodies, particularly fully human monoclonal antibodies, which can be an option for immune passive therapy, have not yet been discovered.

**Methodology/Principal Findings:** This study generated LALA-mutated human monoclonal antibody clone B3B9 (LALA-B3B9 HuMAb) that can neutralize all four DENV serotypes without enhancing viral activity. The number of infected cells obtained with the ADE assay was compared among wild-type antibody (B3B9), and modified Fc, LALA-B3B9 HuMAb, and N297Q (N297Q-B3B9), with or without complement proteins. Moreover, the therapeutic efficacy of these HuMAbs against ADE infection by competing with natural antibodies in patients with acute dengue was determined using the *in vitro* suppression-of-enhancement assay in K562 cells.

The novel Fc-modified antibody LALA-B3B9 (Leu234Ala/Leu235Ala mutations) could have a therapeutic effect. Further, it exhibited neutralization properties against all dengue virus serotypes without triggering ADE activity at any antibody concentration. This outcome was similar to that of the previous Fc-modified N297Q-B3B9 antibody (N297Q mutation). Moreover, the effect of complement protein on enhancing and neutralizing activities was evaluated to assess unwanted inflammatory responses to these therapeutic antibodies.

**Results** showed that the elimination of complement activity could reduce the severity of dengue. The activities of LALA-B3B9 and N297Q-B3B9 HuMAbs were complement-independent in all dengue virus serotypes.

**Conclusions:** LALA-B3B9 and N297Q-B3B9 HuMAbs can prevent the suppression-of-enhancement activity in K562 cells caused by human DENV2. Hence, they are promising candidates for dengue treatment.

**Author Summary:** Dengue is among the most important mosquito-borne infections that cause serious health issues in tropical and subtropical countries. There are no approved effective antiviral therapies available. The significant challenges for developing efficient therapeutic drugs include the presence of multiple serotypes and antibody-dependent enhancement. To overcome these issues, the LALA-mutated human monoclonal antibody clone B3B9 (LALA-B3B9) was generated. This antibody potently neutralizes all dengue virus serotypes and eliminates viral enhancement activity. Moreover, compared with the previous Fc-modified antibody clone N297Q-B3B9, LALA-B3B9 and N297Q-B3B9 HuMAbs do not activate the complement response, which is advantageous in reducing complement-mediated vascular leakage *in vivo*. In addition, they can compete with natural antibodies causing antibody-dependent enhancement in patients with acute dengue. Hence, LALA-B3B9 and N297Q-B3B9 HuMAbs have promising effects against dengue and can be further developed as a therapeutic drug in the future.

## Introduction

Dengue is an arthropod-borne viral disease that is endemic in tropical regions and is a major public health issue. The dengue virus (DENV) is a member of the *Flaviviridae* family. This virus contains a positive-sense RNA genome that encodes three structural proteins (envelope glycoprotein, nucleocapsid protein, and precursor membrane protein) and seven nonstructural proteins (NS1, NS2A, NS2B, NS4A, NS4B, and NS5) [1,2]. The envelope protein is important for both viral attachment and cell fusion. This protein comprises three structural domains (EDI, EDII, and EDIII). A recent study showed that the fusion loops of envelope protein domain II, which is highly conserved in flaviviruses, are the predominant target of cross-neutralizing antibodies in the DENV serotypes. Thus, this epitope is a promising target for developing treatments [3–6].

Dengue is an infection caused by any of the four genetically related but antigenically distinct DENV serotypes (DENV1–4). The symptoms can vary and range from mild or asymptomatic to severe hemorrhage and shock due to the phenomenon called antibody-dependent enhancement (ADE) or ADE activity [7]. ADE occurs by sub-neutralizing antibody–virus immunocomplexes that facilitate virus entry into cells via the Fc gamma receptors (FcγR) and promote virus internalization, which increases viral production. Moreover, the Fc region of sub-neutralizing antibodies can induce ADE via complement activation, which is a major innate immune response and promotes inflammation, thereby contributing to pathogenesis [8–10]. Recent studies have found that the cross-linking of virus– antibody–complement C1q complexes to cell surface C1q receptors leads to increased attachment and enhanced viral entry into cells [11,12]. Due to these reasons, these complement factors cause more severe dengue symptoms. Thus, Fc-FcγR abrogation and complement protein C1q interactions can prevent the unwanted side effects of dengue infection.

Our previous study revealed the glycosylation of human monoclonal antibody against DENV (N297Q-B3B9) [13]. This cross-neutralizing HuMAb is targeted to domain II of the envelope proteins, which is a major target epitope of human antibodies for neutralizing and enhancing activity [14,15]. This antibody exhibits cross-neutralization properties and eliminates ADE activity in all DENV serotypes with NT50, ranging from 12 to 0.125 g/mL [13]. However, a previous study found that N-glycans are essential for not only immune effector functions but also stability under physiological or low temperature conditions, pharmacokinetics, and biodistribution [16–18].

To produce an alternative cross-reactive human monoclonal antibody against all DENV serotypes without enhancing their activity, the site-directed mutagenesis method was used to generate the Fc modification recombinant human monoclonal antibody by double mutation at positions Leu234Ala and Leu235Ala (commonly referred to as LALA mutation), which are residues in the antibody’s lower hinge region, to the heavy chain construct (LALA-B3B9). This modified heavy chain plasmid was co-transfected with light-chain plasmid construct to HEK 293T mammalian cells to produce full IgG. Previous studies have reported that mutations at these two positions can reduce FcγR and complement protein C1q binding [11,19,20].

To investigate the effect of the Fc-modified antibody on the binding of complement protein, the neutralizing and ADE activity of the LALA-modified IgG was evaluated and compared with that of the previously generated aglycosylated HuMAb (N297Q-B3B9) and wild-type HuMAb (B3B9) [21]. The complement-dependent antibody activity was assessed to evaluate the neutralizing and enhancing activities of all HuMAbs. Furthermore, the therapeutic efficacy of Fc-modified antibodies (N297Q-B3B9 and LALA-B3B9) and wild-type antibodies in the human serum was investigated *in vitro*.

## Methods

### Ethical statement

All experimental procedures using human samples were preapproved by the Ethics Committee of the Faculty of Tropical Medicine (FTM), Mahidol University (protocol number: FTM ECF-019-05). All donors provided a written informed consent before enrollment.

### Virus and cells

The DENV1 Mochizuki strain, DENV2 New Guinea C (NGC) strain, DENV3 H87 strain, and DENV4 H241 strain were propagated in C6/36 cells that were cultivated in the Leibovitz L15 medium (Hyclone®, the USA) with 10% fetal bovine serum (FBS) (Hyclone®, the USA) and 0.3% tryptose phosphate broth. Vero cells for the neutralization activity test were maintained in the minimal essential medium (MEM) (GE Healthcare UK Ltd., Buckinghamshire, the UK) with 10% FBS. For the ADE test, K562 cells were cultured in the RPMI 1640 medium (Hyclone®, the USA) supplemented with 10% FBS (Hyclone®, the USA). CHO-K1 cells were used for the stable expression of B3B9, N297Q-B3B9, and LALA-B3B9 antibodies. These cells were cultured in the MEM medium supplemented with 10% FBS and 1% nonessential amino acids (Gibco).

### Generation of LALA-mutated human monoclonal antibody clone B3B9

#### Plasmid construction

The variable genes of the heavy (VH) and light (VL) chains of the B3B9 HuMAb were amplified separately via polymerase chain reaction (PCR) from hybridoma cells [22], and cloned into plasmid containing antibody constant region of pQCX plasmid backbone (Pitaksajjakul et al., 2014; Injampa et al., 2017). Antibody mutations were performed over the Fc portion of the heavy chain plasmid regions at positions 234 and 235 from leucine (L) to alanine (A) (L234A, L235A [LALA]) by site-directed mutagenesis with the In-Fusion Cloning System (In-Fusion® HD Cloning Plus; Clontech Laboratories Inc., Shiga, Japan) based on the manufacturer’s protocol [13].

#### Transient expression of LALA-B3B9 mutation antibody

After sequencing of the mutated heavy chain plasmid for confirmation, the heavy- and light-chain plasmids were transfected to 6.8 × 10^6^ HEK293T cells in a T75 cell culture flask for transient expression. The culture medium was collected and used in immunofluorescence assays to determine the DENV-binding activity. The culture medium containing the secreted LALA-B3B9 recombinant immunoglobulin G (rIgG) was purified using the protein A affinity column (GE Healthcare, Chicago, IL). The eluted fractions from purification were subjected to sodium dodecyl sulfate-polyacrylamide gel electrophoresis to determine the purity of the purified antibody. The LALA-B3B9 rIgG concentration was measured using the BCA protein assay kit (Thermo Scientific, Waltham, MA, the USA).

### Indirect immunofluorescence assay for the DENV-binding activity

Indirect immunofluorescence assay was used to assess the binding activity of the LALA-B3B9 antibodies for all DENV serotypes. Vero cells were mock-infected or infected with DENV at a multiplicity of infection (MOI) of 0.1 in 96-well cell culture plates for 72 h. After fixing the cells with 3.7% formaldehyde in phosphate-buffered saline (PBS) and permeabilized with 0.1% Triton X-100 in PBS, the cells were stained with culture supernatant from transfected cells as primary antibody. After incubation at 37°C for 1 h, Alexa Fluor 488-conjugated anti-human IgG (Invitrogen) (1:1,000) was added as a secondary antibody. The fluorescence can be visualized using a fluorescence microscope (IX71, Olympus).

### Immunoglobulin G subclass determination

The IgG subclass of LALA-B3B9 rIgG was determined via PCR [23]. The gene specific primers of the hinge region between the Fab and Fc parts were used to amplify the hinge region of IgG1, IgG2, IgG3, and IgG4 genes (Table 1). The same reverse (Rv) primer was used for IgG1, IgG2, IgG3, and IgG4. For forward primers, Ig1 and 3 use the same primer for amplification. However, these two isotypes differed in terms of the specific product size due to the length of hinge region. The expected sizes of the PCR products are 211 bp for IgG1, 207 bp for IgG2, 346 bp for IgG3, and 210 bp for IgG4. The PCR mixture comprised 0.5 mg of cDNA, 1.25 mM deoxyribonucleotide triphosphate (dNTP), 1 mM Tris HCl (pH 8.0), 1.25 units of ExTaq DNA polymerase (TAKARA, Shiga, Japan), 5 mM KCl, and 5 mM of each primer with distilled water to a final volume of 25 mL. The amplification was performed by 35 cycles at 94°C for 30 s, 65°C for 30 s and 72°C for 30 s. The PCR products were subjected to agarose gel electrophoresis and visualized by staining with SYBR Safe DNA Gel stain (Invitrogen).

**Table 1.**
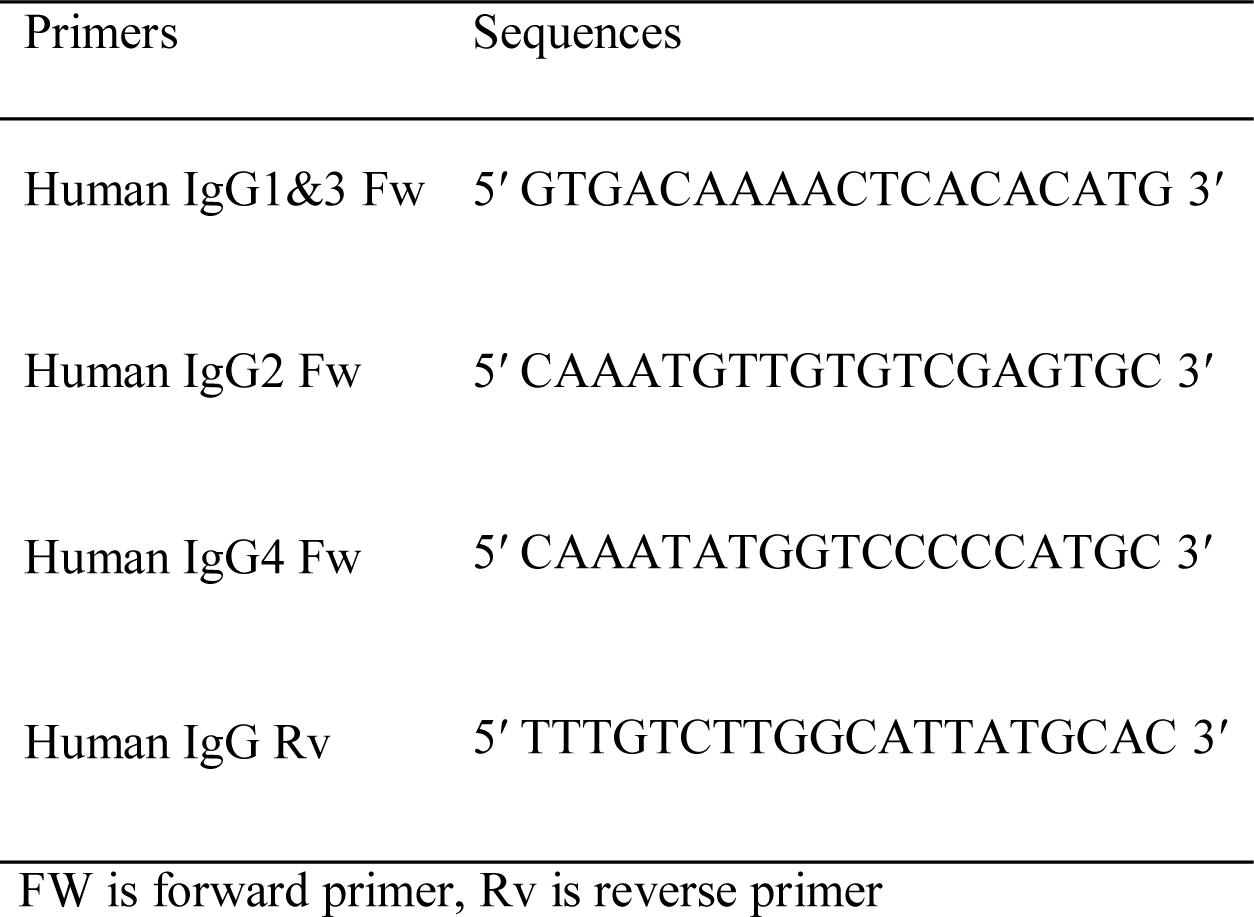
Primers used for IgG subclass determination Primers Sequences.

### Foci reduction neutralization test of the LALA-B3B9 antibodies

The neutralization activity of the LALA-B3B9 antibodies was assessed and compared with that of the previously generated aglycosylated HuMAb (N297Q-B3B9) [13] using Vero cells, as described in a previous study [22]. In brief, DENV at an MOI of 0.01 was preincubated with the respective purified LALA-B3B9 or N297Q-B3B9 HuMAb at 37°C for 1 h before infecting Vero cells in 96-well-cell culture plates. At 2 h post-infection, the plate was overlaid with 2% carboxymethyl cellulose in the MEM medium with 2% FBS. The cells were incubated for 2 days for DENV4 and for 3 days for DENV1, DENV2, and DENV3. Then, the cells were fixed and immunostained using anti-DENV human antibody, followed by Alexa-conjugated antihuman IgG (H+L) (1:1,000 dilution). Neutralization activity was quantified by counting the number of foci for each antibody concentration under a fluorescence microscope compared with the negative control (absence of antibody). The neutralizing antibody titer was considered the minimum IgG concentration yielding a 50% reduction in the focus number (FRNT50).

### Antibody-dependent enhancement assay of LALA-B3B9 antibodies

The ADE activity of the LALA-B3B9 antibodies was performed on FcRIIa-bearing K562 cells, and it was compared with that of N297Q-B3B9 and B3B9 [24]. In total, 36 microliters of serially diluted antibodies were mixed with 50 µl of DENV in 10% FBS RPMI medium in 96-well poly-L-lysine-coated plates (Corning Inc., New York, NY, the USA). The mixture was incubated for 2 h at 37°C, supplied with 5% CO_2_. The mixture was then added with 50 µl K562 cells at a density of 2 × 10^6^ cells/mL and incubated at 37°C under 5% CO_2_ for 2 days. Thereafter, the cells were washed with PBS three times, fixed with an acetone/methanol fixing solution at −20°C, and immunostained with anti-DENV human antibody, followed by Horseradish peroxidase-conjugated antihuman IgG (H+L). The signal was developed with a DAB substrate solution (KPL, Gaithersburg, MD, the USA). The ADE activity was measured by counting virally infected cells manually under a light microscope at a magnification of 10x. Stained cells were counted in three random fields using a 10 × 10 GRID Microscope Eyepiece Micrometer. The total infected cell count in a well was calculated from the average number of stained cells in one grid multiplied by 153.86. The area of one well was 153.86 times greater than grid square. The neutralization and enhancement activity were determined by comparing with the cutoff value, which was calculated by the average number of infected cells without antibody plus the standard deviation (SD) of the percentages of infected cells obtained from the four negative controls. The antibody concentration with a higher and lower number of infected cells compared with the cutoff value was considered ADE and neutralization, respectively.

### Stable expression of B3B9, N297Q-B3B9, and LALA-B3B9 antibodies

The generation of a stable antibody-secreting cell line was described in a previous study [13]. Briefly, CHO-K1 cells were seeded at 1.3 × 10^6^ cells in a six-well-cell culture plate. The heavy and light-chain expressing constructed plasmids, which contained puromycin- and hygromycin-resistant genes, respectively, were co-transfected with transfection reagent (Lipofectamine^®^ 2000) and added to the cells. After 2 days, these cells were incubated in 10% FBS and 1% NEAA MEM containing puromycin and hygromycin at 8 and 800 µg/mL, respectively. After 7 days, all cells in the control well without plasmid died. However, the cells that survived with antibiotic treatment were used for cell cloning by limiting dilution on 96 well-cell culture plates with selection media. The selected single-colony clones were scaled up for further characterization.

### Complement-induced antibody-dependent activity assay

Serial antibody dilutions were incubated with DENV with or without rabbit complement (Cedarlane®, Ontario, Canada) for 2 h at 37°C. The mixture was mixed with 50 µl of 2 × 10^6^ K562 cells/mL and incubated for 2 more days. After fixation and immunostaining, the cutoff between enhancing and non-enhancing activities was calculated using the average percentages of infected cells obtained from the six negative controls plus SD in the same set of experiments [9].

### *In vitro* suppression-of-enhancement assay in K562 cells

#### Serum preparation

A single DENV2 human serum sample collected from patients with dengue infection at the acute phase (days 1–6) was used in this assay. After collecting whole blood samples, the serum was separated via centrifugation, heat-inactivated, and frozen at −80°C.

#### *In vitro* suppression-of-enhancement assay

First, the optimum serum concentration of DENV2 with the greatest enhancement of distinct dengue virus serotypes in K562 cells was determined (1/4000 serum dilution for DENV1 and DENV2, 1/1000 serum dilution for DENV3 and DENV4). Then, the patient serum at the predetermined concentration was mixed with DENV at an MOI of 0.1. The mixture was then incubated for 1 h at 37 °C with 5% CO_2_ prior to the addition of two-fold serially diluted HuMAbs beginning at 1,000 ug/mL. After incubation for 1 h at 37°C, 50 µl of 2 x 10^6^ cells/mL K562 cells were added to the serum–virus–antibody mixture. At 48 h post-infection at 37°C with 5% CO_2_, the cells were fixed by acetone/methanol fixing solution at –20°C, and immunostaining for the viral antigen was performed by incubating with anti-DENV E protein human antibody for overnight at 4°C. After incubation, the plate was washed three times and incubated with Horseradish peroxidase-conjugated antihuman IgG (H+L) diluted in 0.05% Tween-20 and 1% FBS in PBS for 1 h at 37°C. To visualize the infected cells, the signal was developed with a DAB substrate solution (KPL, Gaithersburg, MD, the USA). The average plus the SD of the number of percentages of infected cells obtained with six negative controls set in the same experiment was used as the cutoff value to differentiate neutralizing or enhancing activities [25].

### Statistical analysis

All results were expressed as mean ± SD. All calculations were performed using the GraphPad Prism 6 software.

## Results

### Generation and cross-reactivity of the Fc-modified LALA-B3B9 mutation antibody

The substitution of amino acids at positions 234 and 235 from leucine to alanine in the Fc portion of monoclonal antibody B3B9 was confirmed via DNA sequencing. The cross-reactivity of the LALA-B3B9 antibodies to all DENV serotypes was determined using the immunofluorescence assay and compared with that of the negative control of mock-infected cells. Results showed that LALA-B3B9 antibodies were reactive to all DENV serotypes (Fig 1A). The purity of the produced LALA-B3B9 antibody was assessed via Western blot analysis. Results showed that the protein was observed at the expected molecular weights. The size of full-length IgG with both heavy and light chains is approximately 150 kDa (Fig 1B). This antibody was found to be IgG1 using PCR (Fig 1C).

**Fig 1.**
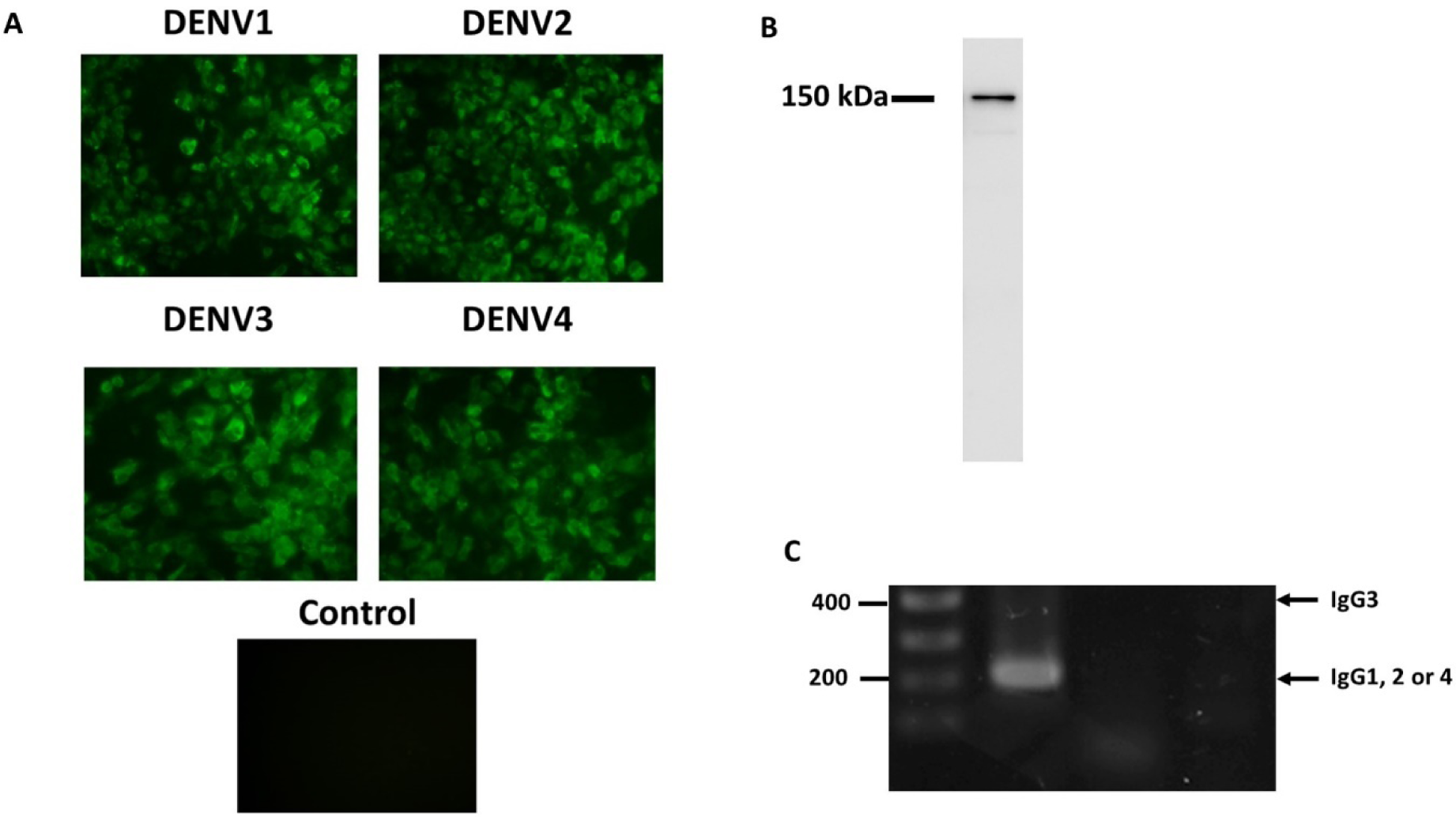
Expression analysis of the LALA-B3B9 antibody. (A) Indirect immunofluorescence assay presented the reactivity of culture supernatants of HEK293T cells transiently expressing target antibody reacted to the envelope protein of four DENV serotypes, with negative control of mock infected cells. (B) Western blot analysis of human IgG from purified culture fluid LALA-B3B9 HuMAb demonstrated the expected band size of full-length IgG. (C) Human IgG subclass determined by PCR technique. The expected sizes of the PCR products are 211 bp for IgG1, 207 bp for IgG2, 346 bp for IgG3, and 210 bp for IgG4. Arrows to the right show the position of each IgG subclass.

### Neutralization of all dengue virus serotypes by the Fc-modified LALA-B3B9 antibody

The FRNT50 test was used to assess the capabilities of LALA-B3B9 mutation HuMAbs to neutralize DENV1–4. The HuMAb neutralizing abilities were compared between the N297Q antibody and the B3B9 antibody, which is the previously modified Fc antibody at position N297Q [13], and the LALA-B3B9 antibody, as described in Fig 2. Both HuMAbs exhibited neutralizing activities against all four DENV serotypes. The LALA-B3B9 antibody strongly neutralized DENV2 and DENV4, and the FRNT50 was < 1 μg/mL. Meanwhile, the FRNT50 concentrations of this antibody against DENV1 and DENV3 were 24.8 and 4.97 g/mL, respectively. The FRNT50 concentrations of N297Q-B3B9 against all serotypes except DENV4 were lower than those of LALA-B3B9 HuMAb. At the highest concentration of 64 μg/mL, the two HuMAbs reduced DENV2 and DENV4 by 100% and DENV3 by nearly 90%. However, these HuMAbs had different neutralizing activities against DENV1. The FRNT50 concentrations of LALA-B3B9 were lower than those of N297Q-B3B9. Table 2 shows the FRNT50 values of the LALA-B3B9 and N297Q-B3B9 antibodies.

**Fig 2.**
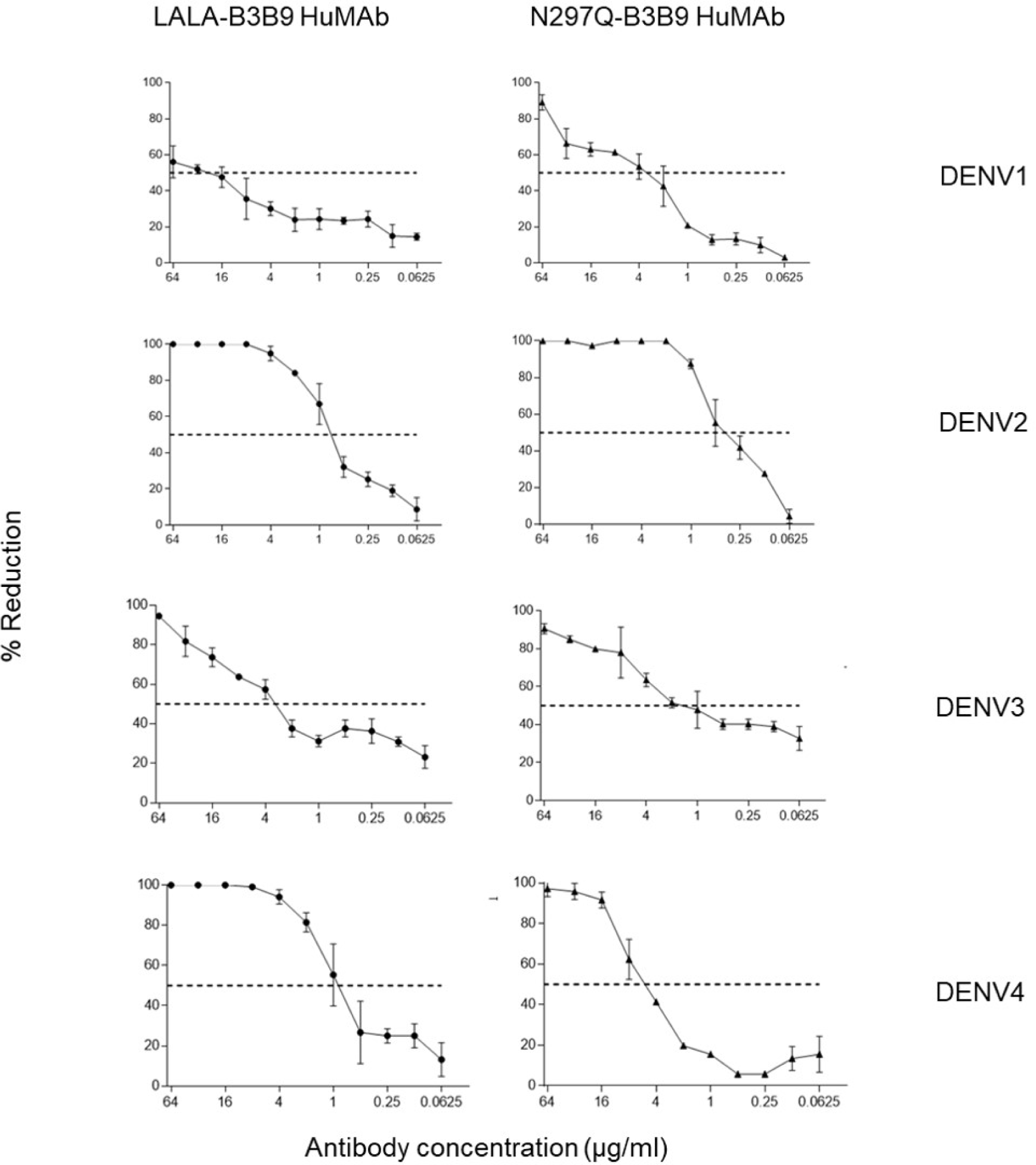
Neutralization activity of LALA-B3B9 and N297Q-B3B9 HuMAbs. Neutralizing activity against DENVs using the average of two independent experiments is shown. Dotted lines indicate 50% neutralizing activity. (The error bars show standard deviation of the experiments)

**Table 2.**
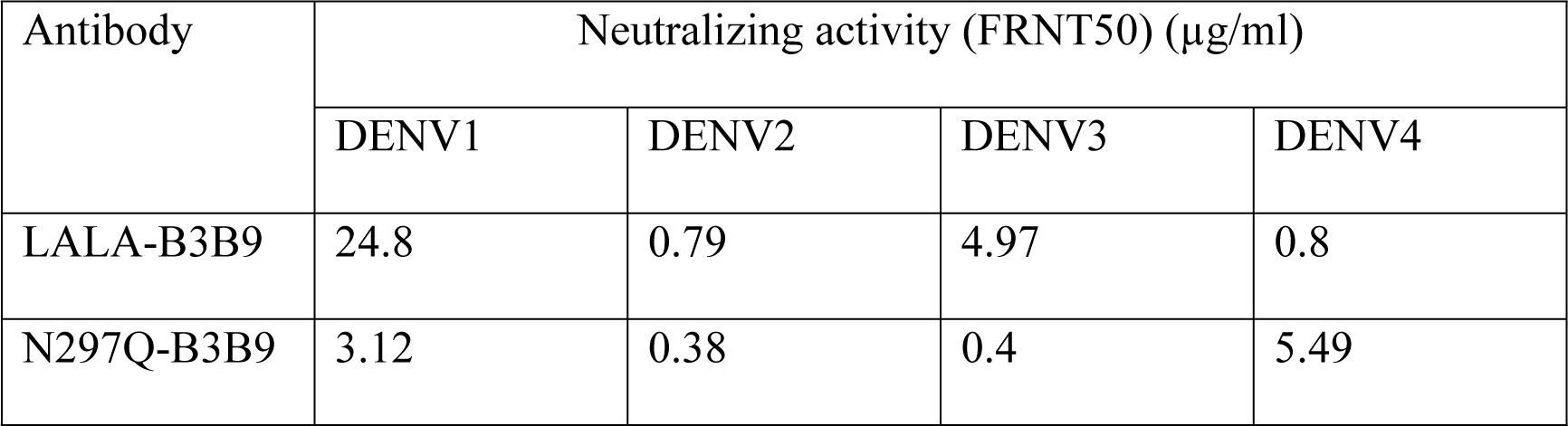
The FRNT50 values comparing LALA-B3B9 and N297Q-B3B9 antibody against all serotypes of dengue virus.

| Antibody | Neutralizing activity (FRNT50) (µg/ml) |  |  |  |
| --- | --- | --- | --- | --- |
|  | DENV1 | DENV2 | DENV3 | DENV4 |
| LALA-B3B9 | 24.8 | 0.79 | 4.97 | 0.8 |
| N297Q-B3B9 | 3.12 | 0.38 | 0.4 | 5.49 |

### ADE activity of the Fc-modified B3B9 LALA mutation antibody

The higher infection rates of immune effector cells occur via the ADE activity, which is mediated by Fc–FcR interactions. Hence, not only neutralizing activity but also ADE activity is a substantial concern for antibody-based therapeutic development. In this study, the ADE activity was determined using FcRII-bearing K562 cells. Fig 3 shows the comparison between wild-type (B3B9) antibody, aglycosylated antibody (N297Q-B3B9), and Fc-modified antibody (LALA-B3B9). The neutralizing and enhancing activities were interpreted by comparing the number of infected cells from each antibody concentration with that of the control without the antibody. Results showed that the ADE activity of the Fc-modified N297Q-B3B9 and LALA-B3B9 antibodies was completely abolished at all antibody concentrations and could neutralize all dengue virus serotypes at higher antibody concentrations, compared with that of the B3B9 antibody with a week neutralizing activity against DENV, particularly DENV1. Further, it activated the ADE activity at sub-neutralizing concentrations in each DENV serotype.

**Fig 3.**
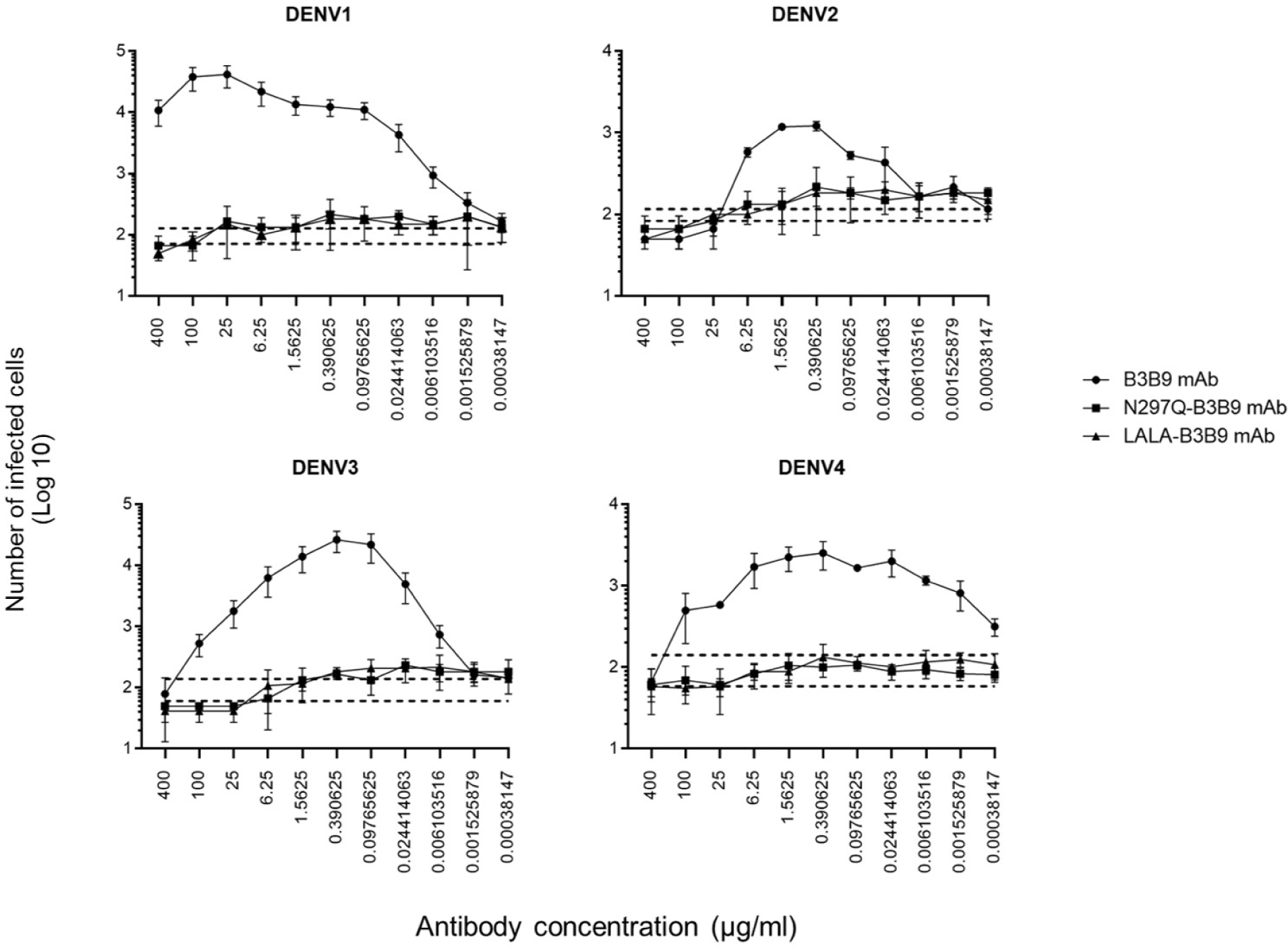
ADE activity of HuMAbs. Neutralization and enhancement activity against the four DENV serotypes in FcRII-bearing K562 cells. The number of infected cells (log10) from each antibody concentration was used to determine the enhancing activity. Dotted lines represent the mean number of infected cells of no antibody control ± SD. Each data point indicates the average result obtained from two independent ADE assays.

### Complement-induced antibody-dependent activity assay

Several studies have found that the excessive consumption of the complement system influences dengue disease severity in humans [9,26]. Therefore, the complement-induced antibody-dependent activity assay was used to determine the balance of neutralization and enhancement in all DENV serotypes with or without complement proteins. Results showed that the Fc-modified N297Q-B3B9 HuMab could neutralize all DENV serotypes without enhancing its infectivity. Moreover, both the neutralizing and enhancing activities of N297Q-B3B9 and LALA-B3B9 HuMab were complement-independent (Fig 4A). The neutralization potency of LALA-B3B9 HuMab is similar with or without complement proteins. LALA-B3B9 HuMab can neutralize all four DENV serotypes and eliminate its enhancing activity (Fig 4B). In contrast, with complement proteins, the ADE activity of B3B9 HuMAb was abolished in DENV1 and DENV2, but still had enhancing activity against DENV3 at antibody concentrations higher than 0.024 g/mL. This antibody showed enhancing activities at all concentrations against DENV4 (Fig 4C), despite the presence of complements, which able to reduce the enhancing activity to a different degree for all dengue virus serotypes.

**Fig 4.**
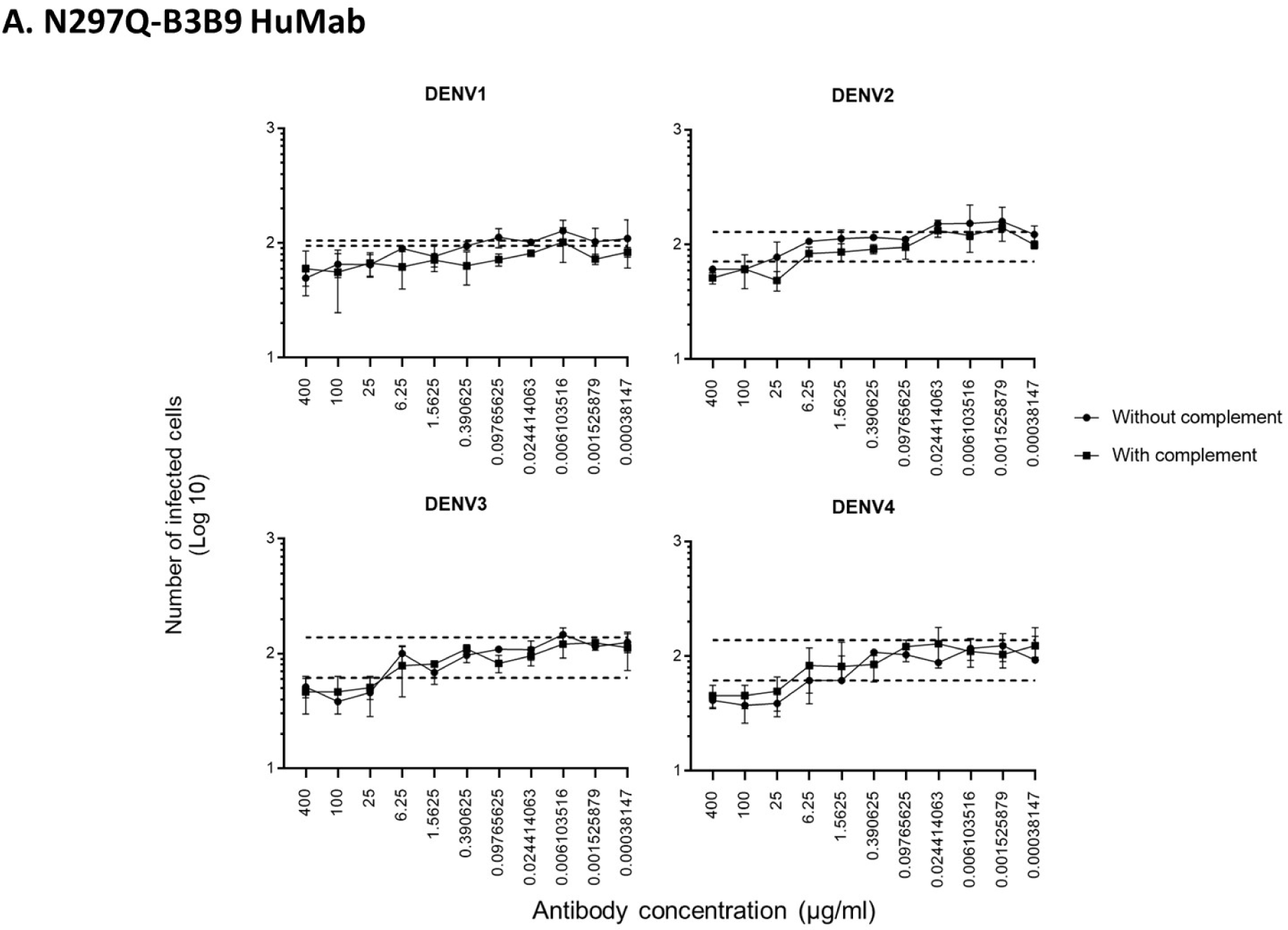
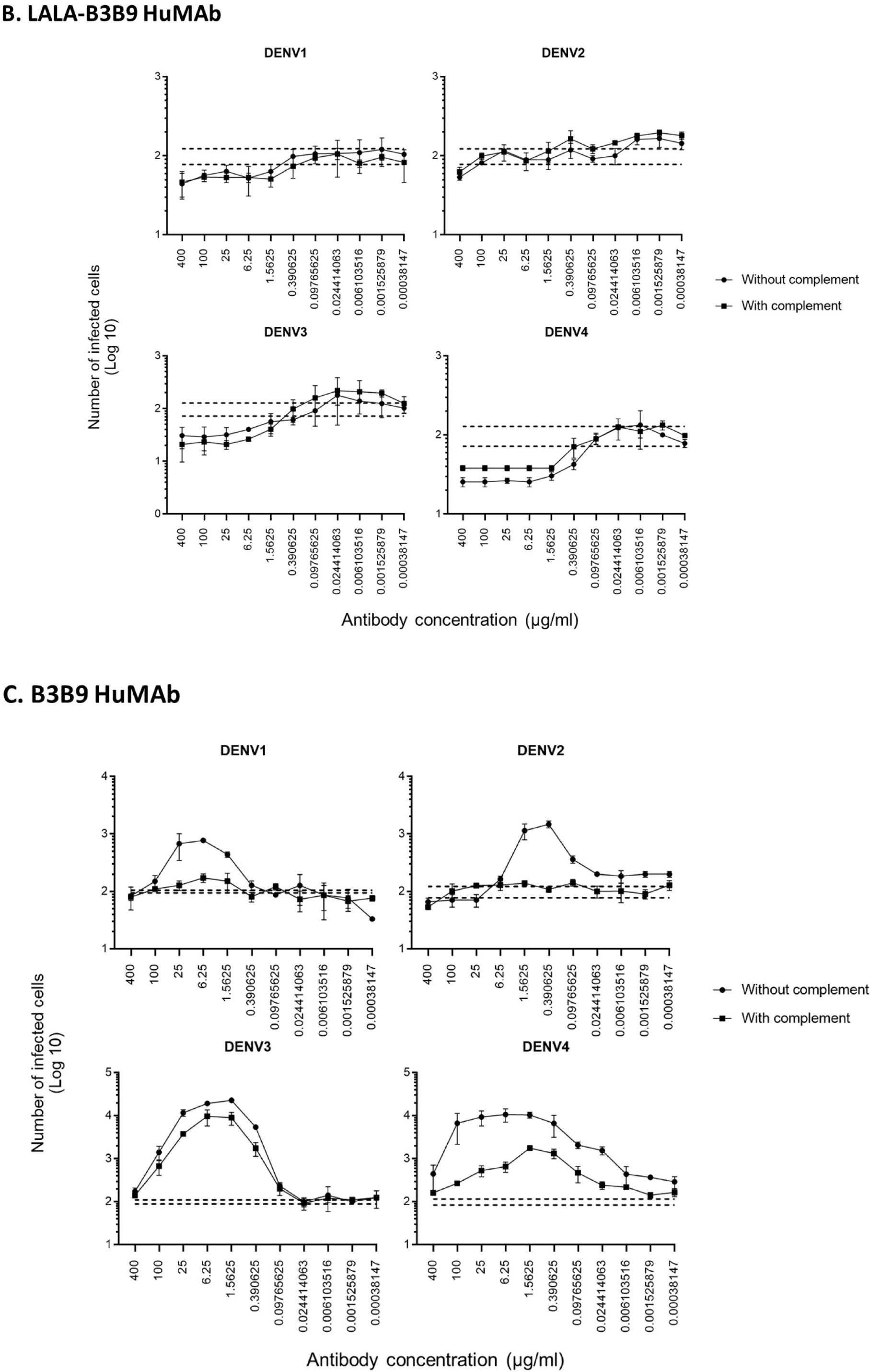
Complement-induced antibody-dependent activity assay of N297Q-B3B9 (A), LALA-B3B9 (B), and B3B9 MAbs (C) against all dengue virus serotypes. Comparison of the number of infected cells from each antibody concentration with and without complement using FcRII-bearing K562 cells. Average of two independent experiments is shown in each data point. Dotted lines display cut-off values for differentiating enhancing and neutralizing activity (the mean number of infected cells of no antibody ± SD). (The error bars show SD of the two repeated experiments.)

**Fig 5.**
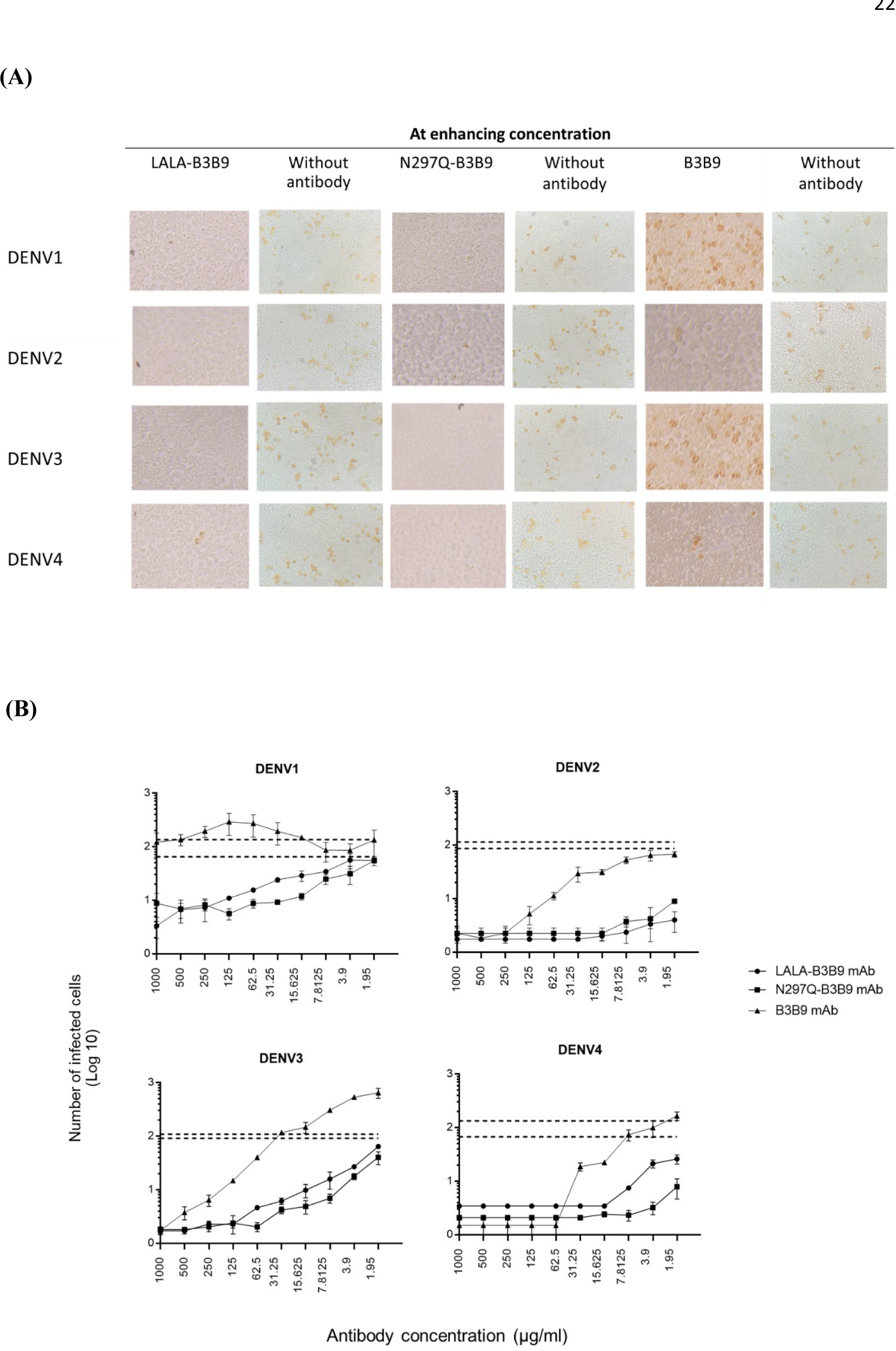
*In vitro* suppression-of-enhancement assay in K562 cells. (A) Comparison of the number of infected K562 cells in enhancing activity (the number of infected cells of mixture of virus, patient serum at enhancing concentration and antibody) and the control (the number of infected cells of mixture of virus and patient serum without antibody). *in vitro* suppression-of-enhancement assay. The peak enhancing titer for DENV2-human immune serum was mixed with dengue virus and each type of antibodies (B3B9, LALA-B3B9 and N297Q-B3B9) at different dilutions and infected to K562 cells. The infection rate was expressed as number of infected cells. Average of two independent experiments is shown in each data point. Dotted lines display cut-off values for differentiating enhancing and neutralizing activity (the mean number of infected cells of mixture of virus and patient serum without antibody ± SD). (The error bars show SD of the two repeated experiments.)

### *In vitro* suppression-of-enhancement assay in K562 cells

The serum samples of patients with acute DENV2 at enhancing concentrations were used to predict the competitive ability of the Fc-modified antibodies and antibodies from natural dengue infection. The use of human serum samples was approved by the Ethical Committees of the Faculty of Tropical Medicine, Mahidol University. The number of infected cells was compared with that of controls (the mean number of cells infected by a mixture of virus and patient serum without antibody ± SD) to determine the neutralizing or enhancing activity. Each generated antibody dilution was mixed with the DENV2 serum at an enhancing concentration that pre-incubated with virus. Then, K562 cells were added. LALA-B3B9 and N297Q-B3B9 had neutralizing activity against all virus serotypes without enhancing their activity. However, B3B9 presented with enhancing activity against DENV1 at antibody concentrations of 500–15.63 g/mL and DENV3 at antibody concentrations of > 31.25 g/mL. However, it could neutralize DENV2 and DENV4 in all antibody dilutions.

## Discussion

Dengue infection is a viral disease that is commonly endemic in subtropical and tropical countries worldwide. This mosquito-borne viral infection is caused by four antigenically distinct dengue viruses (DENV1–4). This infection is a major public health issue in terms of mortality, morbidity, and economic costs [27]. Its symptoms vary, ranging from no symptoms to mild fever to severe disease with vascular leakage, leading to shock and viral hemorrhagic syndrome. The increased risk of severe disease occurs during secondary infection with a virus serotype distinct from that of the prior dengue infection [7,28]. This infection leads to the production of cross-reactive but non-neutralizing antibodies that enhance DENV infection in FcR-bearing cells [29]. To date, there is no specific therapeutic drug against DENV.

The DENV genome encodes three structural proteins (the envelope glycoprotein, nucleocapsid protein, and precursor membrane protein) and seven nonstructural proteins (NS1, NS2A, NS2B, NS4A, NS4B, and NS5). Among several dengue proteins, the envelope protein is a major surface protein that contains most of the neutralizing epitopes and can elicit broad neutralizing antibodies [30–32]. In particular, EDII contains the fusion loop, which is important for viral–host membrane fusion and cell entry [33]. Previous studies have found that antibodies against the E protein of DENV play an important role in the protection and enhancement of the disease, particularly after the primary infection. Moreover, the highly conserved residues at the fusion loop of domain II are the epitopes recognized by the predominantly cross-reactive anti-E antibodies [34]. Hence, these viral protein components of dengue can be targeted effectively to develop antiviral agents.

In a previous study, a fully human monoclonal antibody targeting the fusion loop of envelope protein domain II of the DENV was generated (19). The Fc modification of HuMAb against DENV (N297Q-B3B9) results in the loss of Fc–ligand interactions [13]. This modified antibody can neutralize all DENV serotypes and eliminate the ADE activity in all serotypes in both FcRI- and RII-bearing THP-1 cells and FcRII-bearing K562 cells. However, the removal of glycans at the Asn297 residue, which is the site of conserved N-glycosylation, can cause a substantial change that significantly destabilizes the Fc region and can affect antibody conformation and stability [20,35,36]. Thus, in this study, a novel modified Fc antibody against DENV was established via double mutation at position L234A/L235A (LALA). This was based on previous studies showing that these substitutions reduce binding between the Fc–Fc receptors (FcγRI, FcγRII, and FcγRIII) and the complement component C1q [20]. The neutralizing and enhancing activity of Fc variant L234A/L235A antibody was compared with that of aglycosylated Fc-modified antibody (N297Q-B3B9). Both the Fc-modified N297Q-B3B9 and LALA-B3B9 antibodies can neutralize all DENV serotypes. These two antibodies showed different levels of neutralizing activity. However, ADE was not observed in any antibody concentrations.

The complement system plays an important protective role against infectious diseases. The complement proteins affect the neutralizing/enhancing activities of anti-dengue antibodies [9]. Moreover, the complement was correlated with disease severity in patients with dengue. Failed complement regulatory mechanisms leads to a robust inflammatory response associated with pathological effects, particularly in dengue infection with severe symptoms [10,12]. The mutation at the FC region of the antibody molecules may be involved with the activation of the complement system, which is one of the central parts of innate immunity [37–39], and disease severity in patients with dengue. We then further determined the balance of neutralizing and ADE activity in K562 cells in the presence of complement proteins. For wild-type antibody (B3B9 HuMab), the neutralizing efficacy does not enhance in the presence or absence of the rabbit complement. However, the enhancing activity was reduced, even though not eliminated when complement proteins were applied in each dengue serotype. In contrary, the presence or absence of the complement protein does not mediate the neutralizing or enhancing effects of DENV with N297Q-B3B9 and LALA-B3B9 HuMAbs. These two Fc-modified antibodies had a similar neutralizing activity, and the enhancing activity was inhibited at every antibody concentration. Hence, the modification at position N297Q and Leu234Ala/Leu235Ala can suppress the binding of the Fc portion with FcR and complement proteins.

The IgG1 subclass is commonly induced by DENV infection [40–41]. Hence, this subclass has a high ability to fix complement and bind Fc receptor [42], which is a risk factor resulting in the increased severity of dengue infections. Hence, as our Fc-modified antibodies (N297Q-B3B9 and LALA-B3B9) are IgG1 subclasses, the modification of the Fc region to inhibit binding with the complement and Fc receptor is suitable for further therapeutic application.

Based on our results, the variants of Fc-modified antibodies are considered more advantageous for dengue treatment. To enlighten the application of the modified antibodies during the viremia phase of dengue, acute DENV2-immune serum was used in the enhancement-suppression assay for all dengue virus serotypes for predicting *in vivo* outcomes [24]. The serum concentration with peak enhancement was used in this assay to simulate infection in the serum. By adding dengue serum as an enhancing antibody, the two modified antibodies could compete with the serum antibodies for DENV binding, thereby eliminating serum-derived enhancing activity. This can be possible as the novel Fc-modified antibodies are targeted to the conserved fusion loop of envelope protein domain II, which is also a major epitope recognized by natural human antibody response to DENV [42]. Moreover, these two Fc-modified antibodies showed potent neutralizing activity, and they were selected from several hundred clones of hybridoma derived from antibody-producing cells in patients with dengue [21]. Compared with antibodies targeting other regions, this is the major advantage of anti-EDII antibodies. Thus, N297Q-B3B9 and LALA-B3B9 HuMAbs can prevent enhancement derived in K562 cells from the human DENV2 immune serum and can neutralize viruses in all serotypes.

The novel Fc-modified human monoclonal antibody LALA-B3B9 can neutralize all DENV serotypes without enhancing its activity. This property is similar to that of the previously generated aglycosylated antibody (N297Q-B3B9). Moreover, these two Fc-modified HuMAbs inhibit complement binding activity, which can reduce disease severity [12]. The ability to predict these Fc-modified HuMAb activity in the serum provides novel insight into the mechanism by which modified human monoclonal antibodies can prevent antibody enhancement activity by competing with natural antibodies in the serum of patients with acute dengue.

Overall, considering these promising results, further studies on immune effector functions mediated by Fc–Fc interaction, such as targeted cell elimination via antibody-dependent cellular cytotoxicity and phagocytosis, should be conducted to determine the efficacy of the modified HuMAbs in mediating protection.

## Conclusion

A previous study showed that N297Q-B3B9, an Fc-modified antibody, mutated at the N297Q position of the heavy chain [13]. This antibody can neutralize all DENV serotypes without enhancing their activity. However, changes in the N-linked glycosylation of immunoglobulin can affect effector function causing conformational changes in the Fc domain [17]. To prevent these issues, this study generated LALA-B3B9, a novel Fc-modified antibody targeting the fusion loop on EDII [13]. This antibody has several features that make it appropriate for therapeutic application. That is, it can neutralize all DENV serotypes without enhancing their activity and inhibit the effect of the complement in viral enhancing activity. The effect of the LALA-mutated antibody is similar to that of N297Q-B3B9, a previously generated aglycosylated antibody, without changing the N-glycosylation pattern of the heavy chain. Hence, LALA-B3B9 human monoclonal antibody can be used for dengue treatment in the future.

## Acknowledgments

Authors would like to thank Dr. Nipa Thummasonticharoen who collected dengue patient blood samples and all participants who enrolled in this study.

